# Computational identification of novel Kir6 channel inhibitors

**DOI:** 10.1101/539460

**Authors:** Xingyu Chen, Arthur Garon, Marcus Wieder, Marien J.C. Houtman, Eva-Maria Zangerl-Plessl, Thierry Langer, Marcel A.G. van der Heyden, Anna Stary-Weinzinger

**Affiliations:** Department of Pharmacology and Toxicology, University of Vienna, Vienna, Austria; Department of Pharmaceutical Chemistry, University of Vienna, Vienna, Austria; Department of Medical Physiology, Division of Heart and Lungs, University Medical Center Utrecht, Utrecht, The Netherlands

**Keywords:** KATP channel, Cantú syndrome, molecular dynamics simulation, dynamic pharmacophore, channelopathy

## Abstract

KATP channels consist of four Kir6.x pore–forming subunits and four regulatory sulfonylurea receptor (SUR) subunits. These channels couple the metabolic state of the cell to membrane excitability and play a key role in physiological processes such as insulin secretion in the pancreas, protection of cardiac muscle during ischemia and hypoxic vasodilation of arterial smooth muscle cells. Abnormal channel function resulting from inherited gain or loss-of-function mutations in either the Kir6.x and/or SUR subunits are associated with severe diseases such as neonatal diabetes, congenital hyperinsulinism or Cantú syndrome (CS). CS is an ultra-rare genetic autosomal dominant disorder, caused by dominant gain-of-function mutations in SUR2A or Kir6.1 subunits. No specific pharmacotherapeutic treatment options are currently available for Cantú syndrome. Kir6 specific inhibitors could be beneficial for the development of novel drug therapies for Cantú syndrome, particular for mutations, which lack high affinity for sulfonylurea inhibitor glibenclamide. By applying a combination of computational methods including atomistic MD simulations, free energy calculations and pharmacophore modelling, we identified several novel Kir6.1 inhibitors, which might be possible candidates for drug repurposing. The *in silico* predictions were confirmed using inside/out patch-clamp analysis. Importantly, Cantú mutation C176S in Kir6.1 and S1020P in SUR2A, retained high affinity towards the novel inhibitors. Summarizing, the inhibitors identified in this study might provide a starting point towards developing novel therapies for Cantú disease.

## Introduction

Cantú syndrome is a rare genetic autosomal dominant disorder caused by dominant gain-of-function mutations in the ATP-dependent potassium channel subunits ABCC9 (Harakalova et al., 2012; Van Bon et al., 2012) and KCNJ8 (Brownstein et al., 2013; Cooper et al., 2014, 2017), encoding SUR2 and KIR6.1 respectively. Cantú patients are chronically ill; they suffer from congenital hypertrichosis, distinctive facial features and cardiac defects (Cantú et al., 1982; Nichols et al., 2013; Scurr et al., 2011) and have a decreased life expectancy. Currently, no specific pharmacotherapeutic options are available to treat the disease (Kharade et al., 2016).

Recent breakthroughs in solving atomic and near-atomic resolution structures of eukaryotic inward rectifier potassium channels provide an excellent opportunity to investigate the structural basis of CS mutations and for developing novel therapies for KATP channelopathies. Starting from January 2017, the first (near-)atomic resolution structures (resolution ranges from 3.63 Å to 6.3 Å) of these hetero-octameric complexes have been solved by cryo-EM microscopy by three independent labs (Lee et al., 2017; Li et al., 2017; Martin et al., 2017a, 2017b; Wu et al., 2018). These structures confirm that KATP channels are formed by four Kir6.x pore–forming subunits and four regulatory sulfonylurea receptor (SUR) subunits.

KATP channels couple the metabolic state of the cell to membrane excitability and play a key role in physiological processes such as insulin secretion in the pancreas (Ashcroft, 2005), protection of cardiac muscle during ischemia (Crawford et al., 2002; Nichols and Lederer, 1991; Zingman et al., 2007) and hypoxic vasodilation of arterial smooth muscle cells (Dart and Standen, 1995).

Channel activity is regulated by voltage and ligands. While inhibitory adenosine-triphosphate (ATP) binds to the Kir6.x subunit, magnesium-adenosine-triphosphate-(MgATP) and adenosine-diphosphate-(ADP) activate the channel via interacting with the SUR subunits (MacGregor et al., 2002; Matsuo et al., 1999, 2000; Tanabe et al., 1999; Ueda et al., 1999; Vanoye et al., 2002). Phospholipid phosphatidylinositol-4,5-bisphosphate (PIP_2_), is necessary for channel opening of all inward rectifying potassium channels and binds to the Kir6.x subunit (Huang et al., 1998; Shyng and Nichols, 1998; Zhang et al., 1999). Abnormal channel function, resulting from inherited gain or loss-of-function mutations in either the Kir6.x and/or SUR subunits are associated with severe diseases such as neonatal diabetes, congenital hyperinsulinism and Cantú syndrome (Harakalova et al., 2012; Remedi and Nichols, 2009). Further, SUR2 subunits have been shown to play a role in human neurological disease, including prevalent diseases of the aged brain (Nelson et al., 2015).

Pharmaceutical interventions in KATP channels include sulfonylurea-like inhibitors such as glibenclamide and channel openers, such as diazoxide, which are clinically used to treat neonatal diabetes and hypertension and target the sulfonylurea subunits (Gribble and Reimann, 2003; Pearson et al., 2006). More recently, side effects due to inhibition of KATP channels have been reported as well. For example, in 2011, Yu et al. (Yu et al., 2011) reported that all isoforms of KATP channels are blocked by rosiglitazone (RSG) at micro molar concentrations, which could be harmful due to promotion of adverse cardiovascular effects. RSG is a high-affinity agonist of the peroxisome proliferator-activated receptor γ, which was introduced in 1999 for the treatment of type II diabetes mellitus. The drug increases insulin sensitivity in fat cells by regulating genes involved in glucose and lipid metabolism and might have additional beneficial effects including anti-atherosclerotic, anti-inflammatory and anticancer effects (Brown and Plutzky, 2007). Due to reports of increased risk of myocardial infarction, RSG was withdrawn from the market in Europe in 2010 (Agency European Medicines, 2010) and had its access restricted in the US by the FDA in 2011 (U.S. Food and Drug Administration, 2011). Recently it was shown that this increased cardiovascular risk might be due to modification of different ion channels including Kv4.3, L-type calcium channels and KATP channels (Hancox, 2011; Jeong et al., 2011; Szentandrássy et al., 2011; Yu et al., 2011, 2012). Studies on pigs demonstrate that RSG and other thiazolidinedione drugs can block cardiac sarcolemmal KATP channels *in vivo* at clinically relevant doses (Lu et al., 2008). The reported IC_50_ of this drug is 45 µM for Kir6.2/SURx (pancreatic and heart) channels and 10 μM for vascular Kir6.1/SUR2B. Interestingly, potency has been shown to be even higher in the presence of therapeutic concentrations of sulfonylureas reducing the IC_50_ to 2 μM. Since plasma concentrations of RSG used to treat type II diabetes mellitus are in the range of 3 μM (Cox et al., 2000), block of KATP channels could be a serious problem. Experiments performed on Kir6.2ΔC36 constructs revealed that the drug acts predominantly on the pore-forming Kir6.x subunits and not on the SUR subunits. Further analysis of single KATP channels suggests that the drug suppresses channel activity by extending long-lasting channel closures, most likely via modulating the gating mechanism (Yu et al., 2012).

Kir6 inhibitors such as RSG, which block channels at clinically relevant doses, could provide a good starting point towards development of novel, specific inhibitors, suitable for developing drugs towards treatment of Cantú syndrome. Thus, in this study, we investigated the structural mechanism of pore block of RSG in Kir6 channels. We carried out extensive unbiased full atomistic simulations of drug binding to the closed channel state. Based on the thereby identified binding site, we postulate a structural mechanism by which the drug might prolong the long-closed state of the channel. Further, structure-based pharmacophore models were constructed to enable identification of novel Kir6 inhibitors, which might be useful for future drug development to treat Cantú disease.

## Results and discussion

Based on the experimental finding that RSG predominantly acts on the long-closed state of Kir6.1 channels (Yu et al., 2012), a homology model of the Kir6.1 pore model was constructed using the closed state crystal structure of the Kir3.2 channel (Protein Data Bank (PDB) code: 3SYA (Whorton and MacKinnon, 2011)) as template. A sequence alignment is shown in Supplementary Figure 1A. The root mean square deviation (RMSD) of the Kir6.1 model converged to ~ 4 Å at around 100 ns, indicating that the simulated systems are stable and at equilibrium (see Supplementary Figure 1C). Starting in October 2017, the first atomic resolution structures of the KATP channel formed by Kir6.2 and SUR1 were published (Lee et al., 2017; Li et al., 2017; Martin et al., 2017a, 2017b; Wu et al., 2018). Thus, we compared the structural differences of our Kir3.2 based homology model with the new Kir6.2 templates. Due to the low RMSD of the structural alignments (< 1.5 Å, see Supplementary Figure 1B) we continued to use the Kir6.1 homology model based on the Kir3.2 template in subsequent molecular dynamics (MD) simulations.

**Figure 1.**
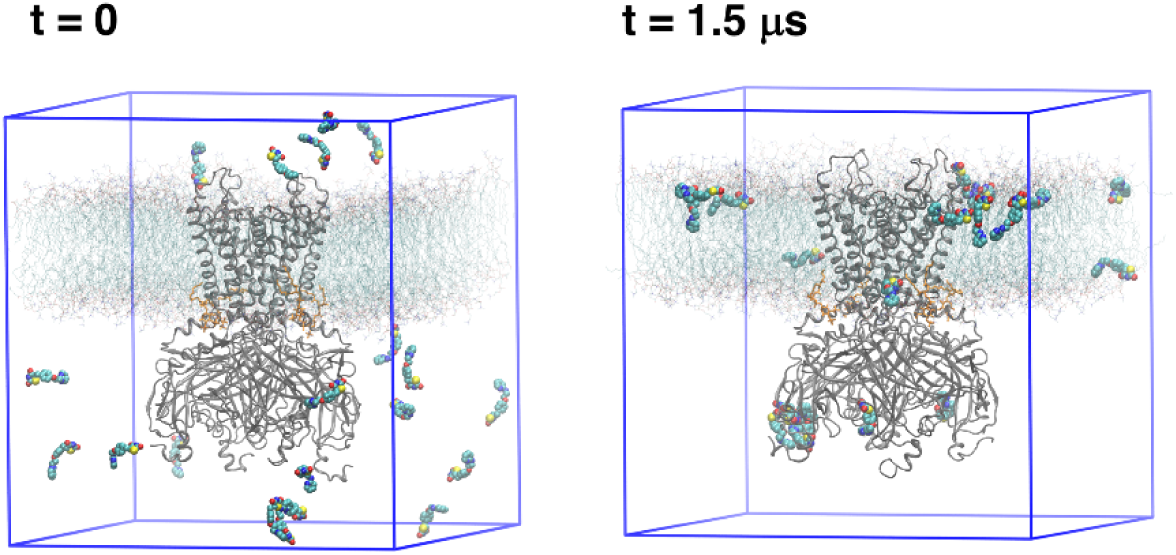
Simulation setups: The first and last frame of free MD simulation. Protein (grey cartoon) is embed in the lipid bilayer (cyan lines with red head groups). As a starting simulation setup (t=0), 20 ligands (shown as spheres) were randomly distributed in the solvent. After a 1.5 μs MD simulation, the ligands either are bound to the protein or entered the lipid phase.

### Unbiased μs time scale MD simulations identify putative RSG binding sites

In an effort to identify the putative binding site of RSG (5-[[4-[2-[methyl(pyridin-2-yl)amino]ethoxy]phenyl]methyl]-1,3-thiazolidine-2,4-dione)), and its main metabolite N-desmethyl Rosiglitazone (5-[[4-[2-(pyridin-2-ylamino)ethoxy]phenyl]methyl]-1,3-thiazolidine-2,4-dione), N-RSG for short, full atomistic molecular dynamics simulations were performed. Specifically, the binding was probed by adding 20 molecules (10 x S conformer, 10 x R conformer, since the prescribed drug is a racemic mixture) randomly into the solvent, leading to an effective drug concentration of ~ 170 mM. As seen in Figure 1, 13 ligands partition into the lipid membrane within 1,5 μs. Three out of 20 molecules occupied sites close to the protein for 1.3 – 1.5 μs (see Table 1). The three major binding sites (Figure 2) identified are: close to the PIP_2_ binding site, denoted site A; the interface between two cytoplasmic domains (CTDs), denoted site B, and between the β-sheet βD and the βG-αG loop, denoted site C.

**Table 1.** Three binding sites observed during unbiased MD simulations and their corresponding occupation time. All three ligands occupied the binding site for at least 1 μs and until the end of the simulation. Ligands migrate to binding site A at 300 ns, to binding site B right after the equilibration, and to binding site C at 200 ns.

| Binding site | Binding time of ligand to binding site | Stereochemistry of ligand |
| --- | --- | --- |
| A | 1.2 μs (300 ns - 1.5 μs) | R conformer |
| B | 1.5 μs | S conformer |
| C | 1.3 μs (200 ns – 1.5 μs) | S conformer |

**Figure 2.**
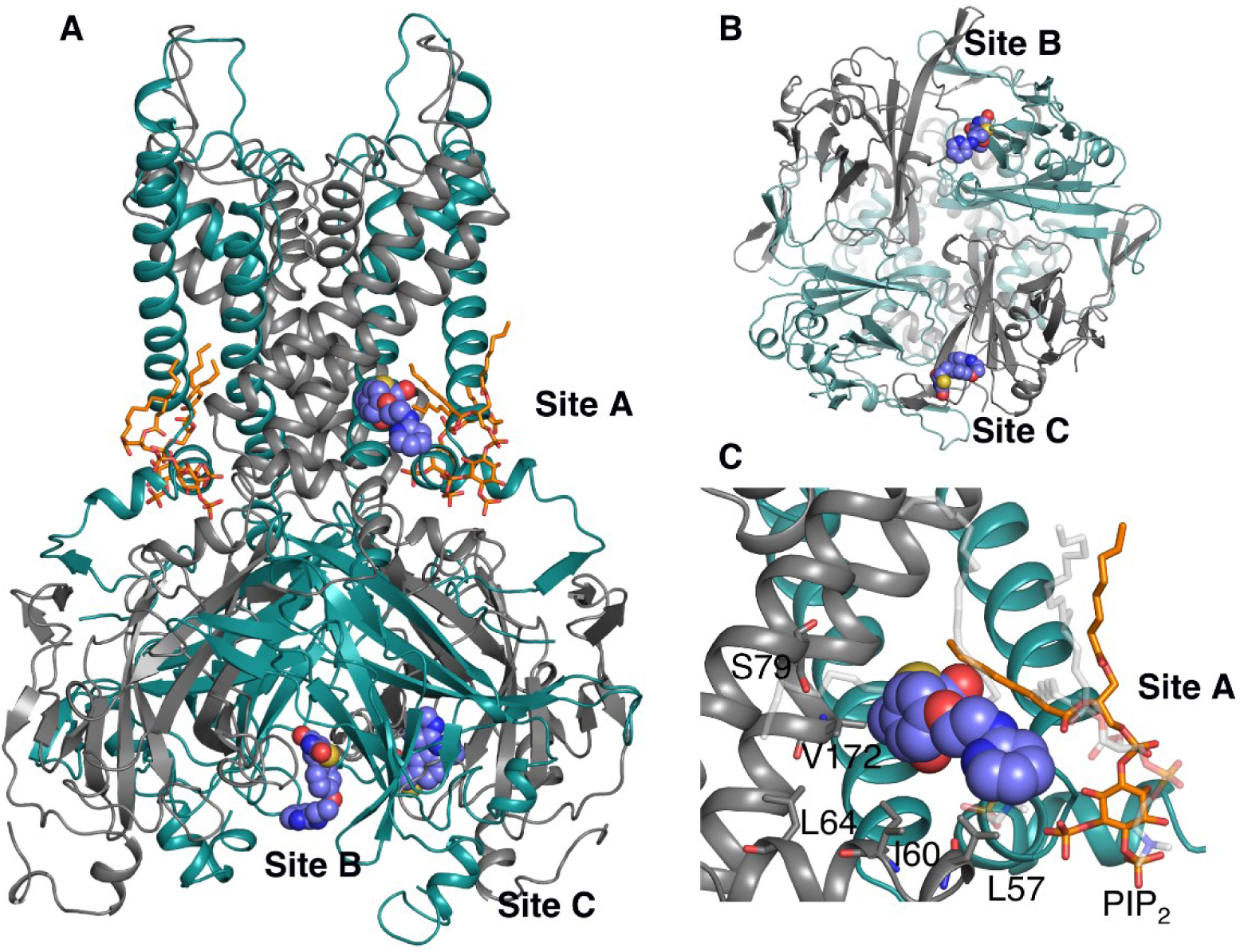
Three binding sites observed after 1.5 μs unbiased MD simulation. The protein is represented as cartoon; adjacent subunits are colored in green and grey, respectively. PIP_2_ molecules are shown as orange sticks. Ligands are represented as purple spheres. **(A)** Side view of the whole protein highlighting the three ligands bound to binding sites A – C. Site A: in close proximity to the PIP_2_ binding site; Site B: at the interface between 2 CTDs; Site C: between the β-sheet βD and the βG-αG loop. **(B)** Bottom view of Site B and Site C. **(C)** Detail view of site A: residues within 6 Å of the ligand are labeled and presented as grey sticks. The ligand mainly forms hydrophobic interaction with LEU57 and ILE60; additionally, it forms hydrophobic and hydrogen bond interactions with lipid molecules (transparent sticks in light grey).

Binding site A is of particular interest since the ligand binds close to the transmembrane gate and the PIP_2_ binding site. Binding includes mainly hydrophobic interactions with residue LEU57 and ILE60. Additionally, hydrophobic and hydrogen bond interactions with POPC lipid molecules are observed. Even though PIP_2_ is within 5 Å of ligand no specific interactions with RSG are observed. All residues within 6 Å of ligand are shown in Figure 2C. Given the importance of this area for channel gating (Zhang et al., 2015), we decided to perform a more exhaustive sampling of this region. We therefore used computationally less demanding docking simulations with both compounds, followed by 250 ns MD simulations of the best-scored pose for RSG, and 200 ns of the best-scored pose for N-RSG. This way, strengthened binding interactions (defined as increased accessible surface area: 0.9 nm^2^ for site A vs. 3 nm^2^ when using the best docking pose) were obtained. We refer to this new site A as site A_ref.

To further characterize the three binding sites, the potential of mean force (PMF) of the ligands at site A_ref, B and C, were calculated using umbrella sampling (US) simulations. As can be seen in Figure 3, the PMFs reveal clear differences between the three sites. Binding of RSG to site A_ref is most favorable. Since PMF calculations reveal shallow binding for ligands bound at sites B and C, we did not further investigate these two binding sites. Nevertheless, ligand binding site B, might be potentially interesting, since intersubunit interactions in this region have been shown previously to be important for the inactivation process in Kir6.2 channels (Borschel et al., 2017; Lin et al., 2003).

**Figure 3.**
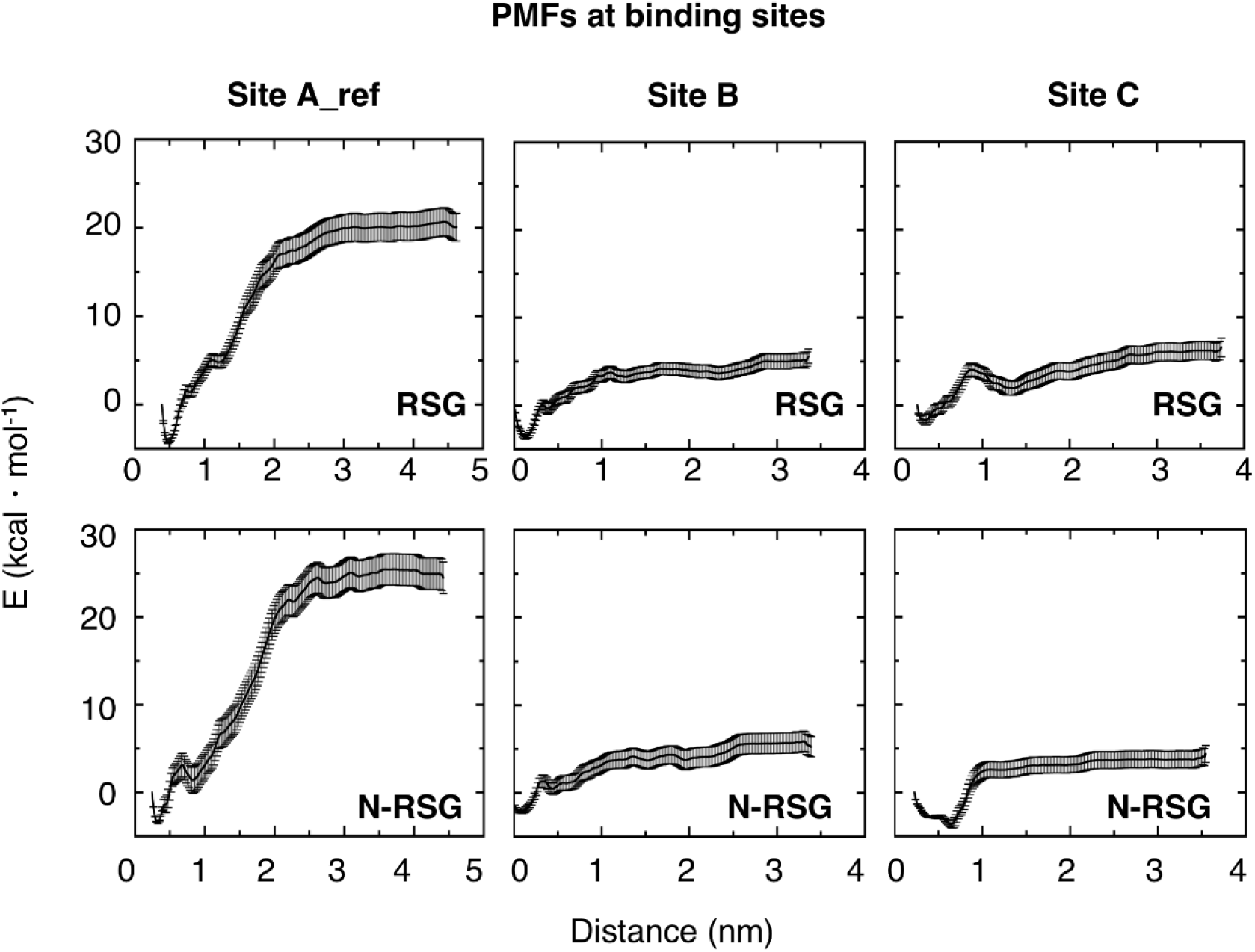
Potential of Mean Forces (PMFs) derived from umbrella sampling for both RSG and N-RSG at the three binding sites. The energy profiles are depicted including their standard deviations. The WHAM histograms are shown in Supplementary Figure 4. Site A_ref shows higher binding affinity for both, RSG and N-RSG, compared to the other binding sites.

### Detailed characterization of binding site A_ref

The interactions of RSG and N-RSG at site A_ref over 200 ns of the MD simulations were quantified using interaction matrices as described in the methods section. Ligands were decomposed as five parts (Figure 4A and Supplementary Figure 2): the pyridine (Ring A), the benzene (Ring B), the thiazolidinedione (Ring C), the linker connecting pyridine and benzene (Linker D), and the linker connecting benzene and thiazolidinedione (Linker E). RSG mainly formed hydrophobic interactions with the binding site throughout the trajectory. More than 75% of the frames contain hydrophobic interactions between the Ring A and residue PHE76, VAL172 and MET173, and between the Linker E and TRP69. Residues LEU64, ILE75, PHE80, ILE168 and ILE169 also frequently formed hydrophobic interactions with RSG (Figure 4B). The interaction map of N-RSG molecule reveals a similar trend indicating mainly hydrophobic interactions with the binding site (see Supplementary Figure 2). Both ligands also form hydrogen bonds with POPC lipid molecules, which are not included in the interaction map calculations.

**Figure 4.**
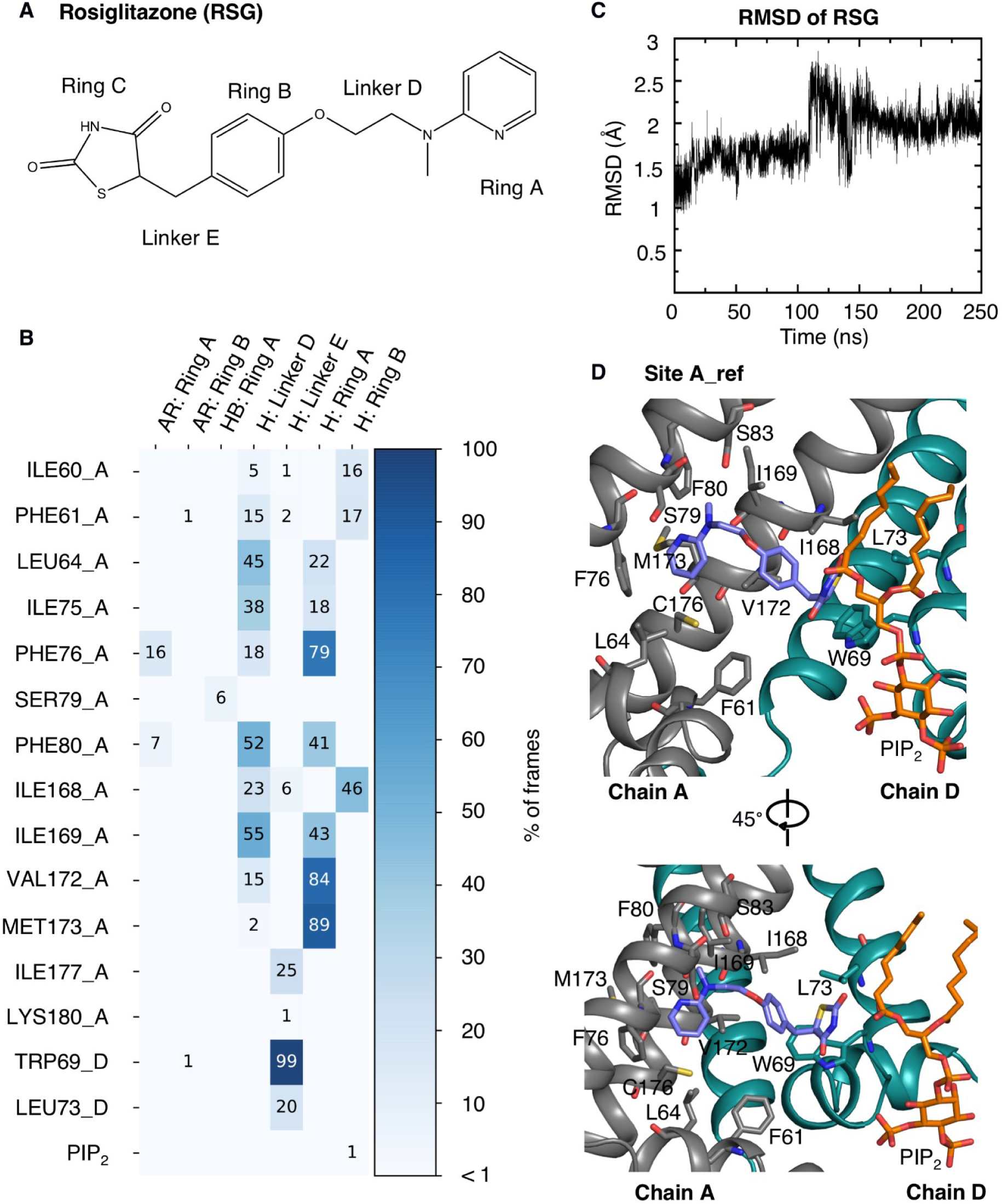
RSG interactions with the protein and PIP_2_ at binding site A_ref. **(A)** Molecular structure of RSG including the denotation corresponding to the interaction map. **(B)** Interaction map of RSG with protein and PIP_2_ during 200 ns MD simulation. The matrix is colored and numbered by the percentage of frames, in which interactions were observed: aromatic (AR), hydrophobic (H), hydrogen bond (HB). The residues in the Kir6.1 are named by the corresponding amino acid with its residue number and chain ID (A or D). **(C)** RMSD plot of RSG at binding site A_ref during a 250 ns MD simulation. **(D)** Best PMF energy pose: Kir6.1 is represented as cartoon with two neighboring subunits colored in grey and green, respectively. RSG (purple), the surrounding residues within 3.5 Å, and the PIP_2_ (orange) are shown as stick.

### Structure based dynamic pharmacophore models of RSG binding to site A_ref

Understanding the inhibition mechanisms of the pore-forming Kir6.1 subunit could be a first step to develop novel blockers for the treatment of rare disease causing mutations (e.g. Cantú mutations V64M), which are not amenable for sulfonylurea therapy (Cooper et al., 2017). In line with this reasoning we constructed dynamic structure-based pharmacophore models and screened for hits in *DrugBank* (Law et al., 2014), which contains all marketed drugs, by using the common hits approach (CHA) (Wieder et al., 2017). Structure-based pharmacophore models were generated using 5,000 frames from the MD simulation, which included the lipid bilayer but omitted solvent molecules. Pharmacophore models, which contain common pharmacophore features and identical involved ligand atoms, are considered as one representative pharmacophore model. Five representative pharmacophore models (Model 1 – 5) were observed from the frames (shown in Supplementary Figure 3). Model 1 was observed most frequently (> 95%, 4,776 times out of 5,000 of frames). The other four models appeared in less than 5% of the frames. All models share two hydrophobic features formed with pyridine and benzene in the ligand. Model 1 contains one additional hydrogen bond donor with the NH moiety of thiazolidinedione. Model 2 only contains the shared hydrophobic features. Model 3 and Model 4 also share the same hydrophobic features plus one hydrogen bond donor and one hydrogen bond acceptor. Model 5 comprises the shared hydrophobic features and one hydrogen bond acceptor. All five pharmacophore models were used to screen *DrugBank*. The top ranked hitlist for binding site A_ref is shown in Table 2.

**Table 2.** Top ranked hit-list for binding site A_ref. The hit-list was established by screening the dynamic pharmacophore models to the *Drugbank* database. Top 20 approved drugs ranked by CHA score are proposed in the displayed hit-list.

| DrugBank ID | Generic name | CHA score | Number of active pharmacophores |
| --- | --- | --- | --- |
| DB08907 | Canagliflozin | 2.7417 | 3/5 |
| DB01095 | Fluvastatin | 2.7246 | 3/5 |
| DB09351 | Levobetaxolol | 2.7035 | 3/5 |
| DB00917 | Dinoprostone | 1.8892 | 2/5 |
| DB00195 | Betaxolol | 1.862 | 2/5 |
| DB00841 | Dobutamine | 1.852 | 2/5 |
| DB00287 | Travoprost | 1.8516 | 2/5 |
| DB00938 | Salmeterol | 1.8475 | 2/5 |
| DB00179 | Masoprocol | 1.8458 | 2/5 |
| DB00867 | Ritodrine | 1.8104 | 2/5 |
| DB09198 | Lobeglitazone | 1.8102 | 2/5 |
| DB04855 | Dronedarone | 1.8096 | 2/5 |
| DB06817 | Raltegravir | 1.805 | 2/5 |
| DB09570 | Ixazomib | 1.8038 | 2/5 |
| DB01346 | Quinidine barbiturate | 1.8019 | 2/5 |
| DB00204 | Dofetilide | 1.7997 | 2/5 |
| DB01240 | Epoprostenol | 1.7992 | 2/5 |
| DB00662 | Trimethobenzamide | 1.7935 | 2/5 |
| DB08875 | Cabozantinib | 1.7838 | 2/5 |
| DB09330 | Osimertinib | 1.7275 | 2/5 |

### Inhibition of Kir6.2/SUR2A by Travoprost, Betaxolol and Ritodrine

From earlier work it was established that in the absence of pharmacological activation, Kir6.1/Sur2a channels yield very low current amplitude which hampers efficacy assessment of blockers (Cooper et al., 2014). Therefore, Travoprost, Betaxolol and Ritodrine (three top ranked hits (Table 2)) were tested for Kir6.2/SUR2A inhibition instead, using the inside/out mode on HEK293T cell derived excised membrane patches. Travoprost (IC_50_outward=2.46±0.52 μM; Hill coefficient 0.71; IC_50_inward=2.30±1.26 μM; Hill coefficient 0.65) dose-dependently inhibited inward and outward current components of I_Kir6.2/SUR2A_ whereas Betaxolol (IC_50_outward=22.06±2.47 μM; Hill coefficient 0.89) and Ritodrine (IC_50_outward=7.09±0.45 μM; Hill coefficient 0.86) markedly and dose-dependently inhibited the outward component (Figure 5). Betaxolol and Ritodrine induced rectification behavior of the channel, i.e. outward current was more strongly inhibited than inward current (Figure 5B). This induction of rectification is in contrast with inhibition characteristics of Travoprost and RSG. Similar findings have been made with Pentamidine and Pentamidine-Analogue-6 (De Boer et al., 2010; Takanari et al., 2013). These structurally related compounds bind to the same site in the Kir2.1 channel, but whereas Pentamidine induces channel rectification (De Boer et al., 2010), Pentamidine-Analogue-6 inhibited both inward and outward current with similar efficacy (Takanari et al., 2013).

**Figure 5.**
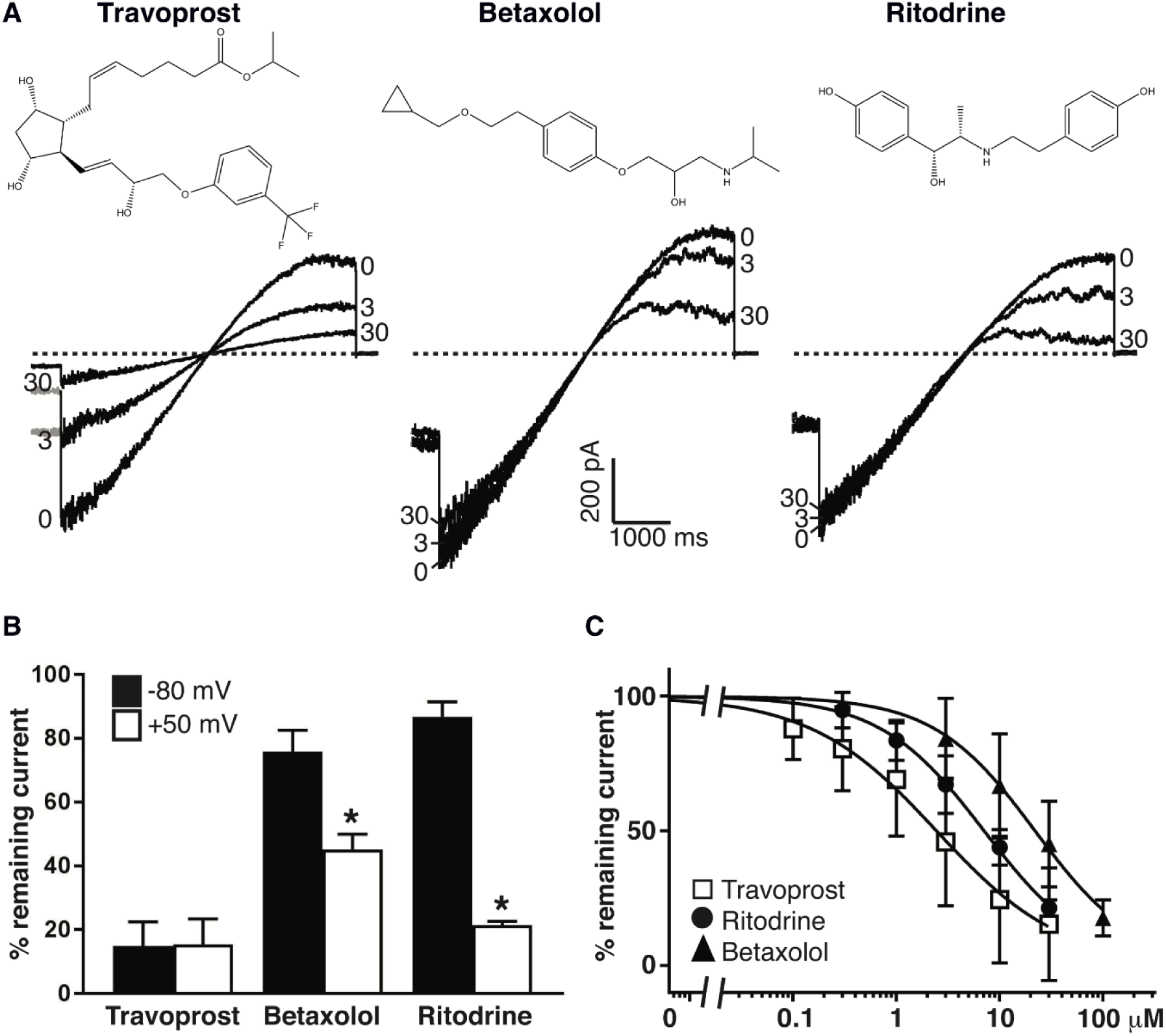
Inhibition of Kir6.2/SUR2A carried I_KATP_ by Travoprost, Betaxolol and Ritodrine. **(A)** Current traces of Kir6.2/SUR2A channels in the inside-out orientation exposed to Travoprost, Betaxolol or Ritodrine at the indicated drug concentrations (0, 3 and 30 μM). Dotted horizontal line at 0 pA. **(B)** Normalized block of inward (black bars, at −80 mV) and outward (open bars, at +50 mV) currents with 30 μM of the indicated drug. *P<0.001 (paired T-test, inward vs. outward; n=8, n=11 and n=9 for Travoprost, Betaxolol and Ritodrine, respectively. **(C)** IC_50_ curves of outward components of Kir6.2/SUR2A in response to different concentrations of Travoprost (open squares; n=8), Betaxolol (black triangles; n=11 or n=7 (100 μM)) and Ritodrine (black circles; n=9 (control and 1 μM), n=8 (0.3 μM) or n=7 (3, 10, 30 μM)). Data were fitted with Hill equation to estimate the IC_50_ values. Data in panels b and c are shown as mean±SEM.

### Further experimental support of RSG binding to site A_ref

Experimental data from Yu et al., 2012 (Yu et al., 2012) revealed that RSG binding to inward rectifier K^+^ channels is Kir6.x specific. Electrophysiology measurements on Kir1.1, Kir2.1 and Kir4.1 channels showed that these channels are insensitive to RSG. Thus, we performed a multiple sequence alignment of these channels and evaluated the conservation of the predicted binding site residues. As shown in Figure 6, differences between Kir6.x and members of other Kir channels at site A_ref can be found at positions 64 (LEU vs. VAL/CYS/PHE), 79 (SER vs. ALA/THR), 172 (VAL/ILE vs. PHE) and 176 (CYS vs. ALA/THR). Of particular interest is position 172, which contains a bulky phenylalanine side chain in non-Kir6.x channels, which would prevent RSG from binding in a similar mode in these channels. Experimental mapping of the binding site has not been performed, possibly due to challenges measuring Kir6.1 subunits (Cooper et al., 2014). Nevertheless, mutational data on the closely related Kir6.2 channel supports that residues in the binding area are critical for normal channel gating (Zhang et al., 2015). Interestingly, two Cantú disease causing gain-of-function mutations (V65M, C176S) (Brownstein et al., 2013; Cooper et al., 2014) are in close proximity to binding site A_ref, further supporting the importance of this region for gating. A further, indirect validation of the binding site was gained by correctly identifying novel hits in *DrugBank*, based on the *in silico* predicted structure-based drug-protein interactions.

**Figure 6.**
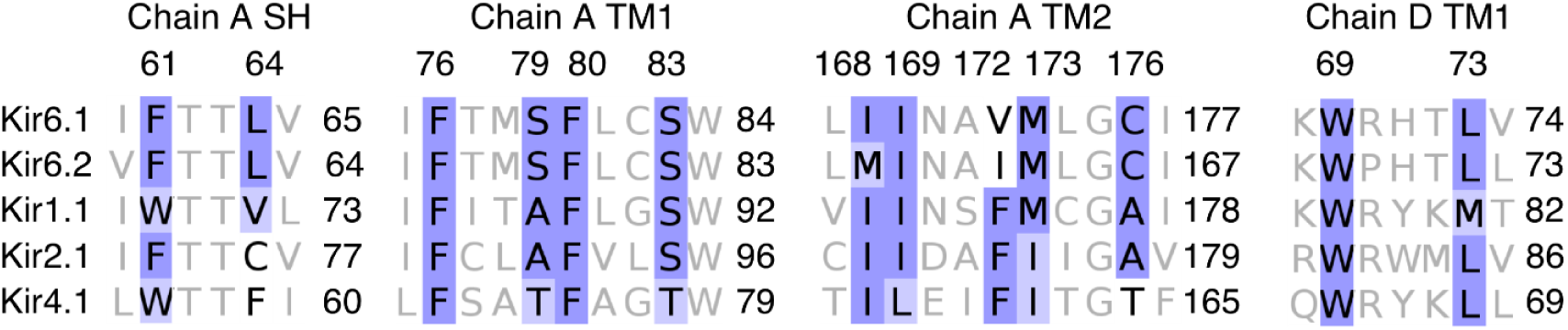
Sequence alignment of Kir6.x, Kir1.1, Kir2.1 and Kir4.1 from binding site A_ref. The alignment of residues on chain A and chain D within 3.5 Å of RSG at the lowest PMF energy pose. Coloring of the alignment was performed using the BLOSUM62 algorithm. SH: slide helix; TM1: transmembrane helix 1; TM2: transmembrane helix 2.

### Suggestions for experimental validation of the binding site

Our modelling predictions suggest that mutating position VAL172 to PHE should decrease or prevent binding of RSG and its main metabolite N-RSG. Unfortunately, mutating the equivalent position in Kir6.2 (ILE162) to PHE does not produce functional channels (Piao et al., 2001), preventing experimental validation of this prediction. Since the binding site is very close to the PIP_2_ molecule, interference of PIP_2_ interactions with the channel are likely. Previous studies on other Kir channels, support drug PIP_2_ interference for drugs such as carvedilol and ivermectin (Chen et al., 2017; Ferrer et al., 2011; Kikuta et al., 2006) or the anti-cancer agent gamboic acid (Scherer et al., 2017). For a recent review see (Heyden et al., 2013).

### Proposed mechanism of drug action

RSG binds at the interface between two subunits, very close to PIP_2_, an essential gating modulator of inward rectifier channels. It is conceivable that the drug interferes with normal channel activation, possibly via “blocking” the activation gate and/or via hindering normal lipid modulation of channels. In line with this reasoning, we observed frequent hydrogen bonds to lipid molecules in our simulations. Further simulations, using different lipid types will be necessary in the future to investigate this possibility. In addition, the predicted binding site A_ref is in close proximity of two previously identified gain-of-function mutations causing Cantú disease (V65M and C176S (Brownstein et al., 2013; Cooper et al., 2014)). So far 100 ns MD simulations on these two mutations have been performed (Cooper et al., 2017), revealing no changes in this region. Thus, in order to investigate if RSG or the newly identified inhibitors might be able to counterbalance the gating disturbance effects of Cantú mutations, we determined dose-response effects of currents mediated by C166S Kir6.2, which is homologues to C176S in Kir6.1.

### Cantú mutations C166S (Kir6.2) and S1020P (SUR2A) are inhibited by RSG, Ritodrine, Travoprost and Betaxolol

Since CS C176S mutant is in close proximity of the predicted binding site A_ref, we performed inside/out measurements on Kir6.2 C166S, the homologues mutation of Kir6.1 C176S, with RSG and the newly identified drugs Travoprost, Betaxolol and Ritodrine. As shown in Figure 7 all drugs dose-dependent inhibit outward current, having IC_50_ similar as WT channels (RSG: WT 25.98±1.49 μM vs. C166S 34.88±2.34 μM n.s.; Ritodrine: WT 7.09±0.45 μM vs. C166S 10.42±0.87 n.s.; Betaxolol: WT 22.06±2.47 vs. 41.16±2.89 μM n.s.) except for Travoprost (WT 2.46±0.52 μM vs. 14.82±2.16 μM, p<0.05).

**Figure 7.**
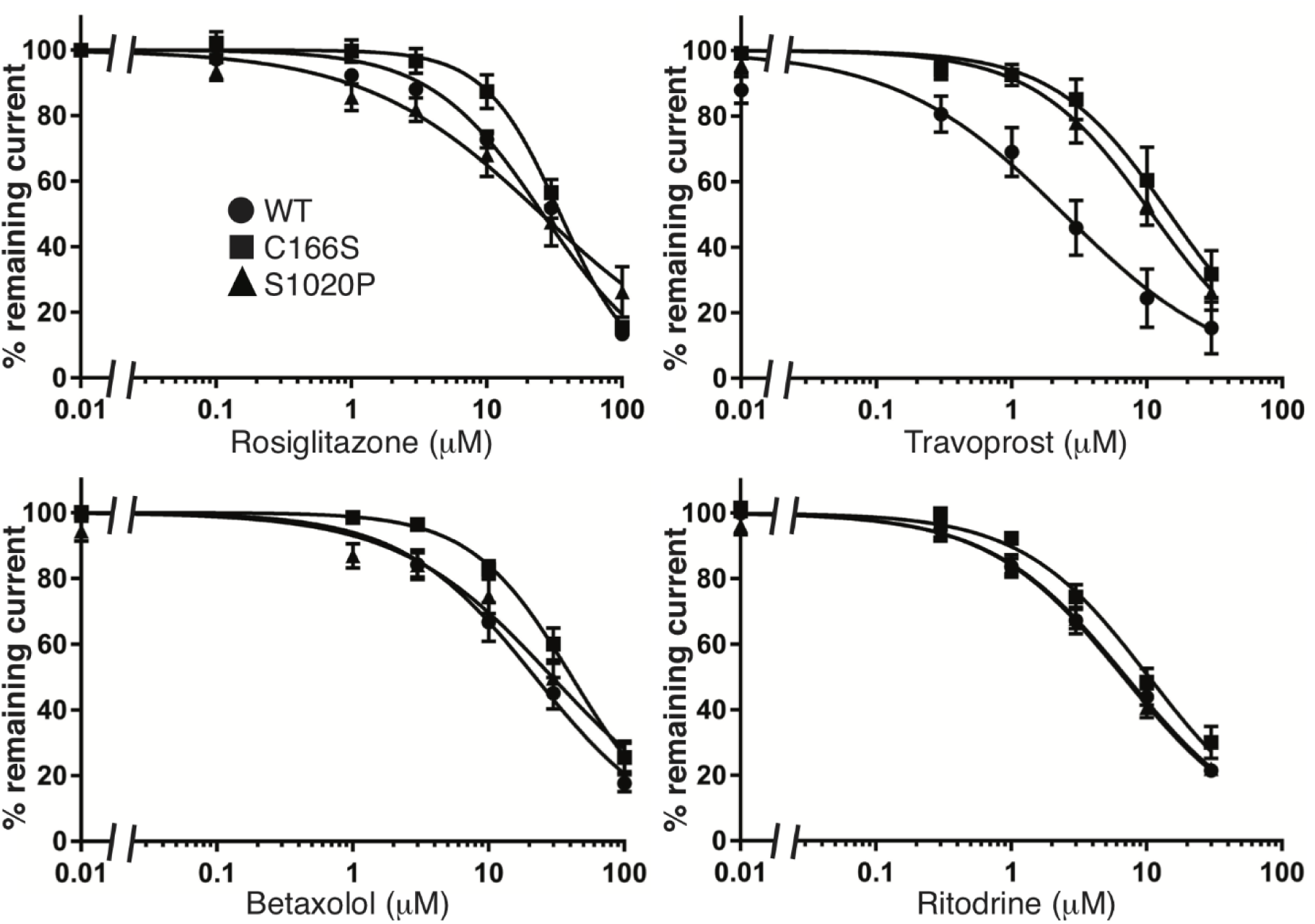
Inhibition of C166S (Kir6.2) and S1020P (SUR2A) by Rosiglitazone, Travoprost, Betaxolol and Ritodrine. IC_50_ curves of outward components of Kir6.2/SUR2A (WT, C166S Kir6.2, S1020P SUR2A) in response to different concentrations of Rosiglitazone, Travoprost, Betaxolol and Ritodrine. N-values are: WT, C166S and S1020P respectively: Rosiglitazone n=8, 8, 6; Travoprost n=8, 7, 11; Betaxolol n=11, 7, 7; Ritodrine n=9, 8, 7. Data were fitted with Hill equation to estimate the IC_50_ values. Data are shown as mean±SEM.

Since the majority of Cantú mutations have been identified in the SUR2A subunits, we also tested the inhibitors on currents mediated by the S1020P SUR2A mutation. Again, mutant channels were sensitive for all for compounds with outward current IC_50_ values (Rosiglitazone: 25.38±4.24 μM; Ritodrine: 6.77±0.49 μM; Betaxolol: 29.51±3.78 μM) not significantly different from WT channels except for Travoprost (10.99±1.28, p<0.05 vs. WT).

## Conclusions

Understanding the molecular mechanisms of inhibition of Kir6.x channels is critical to in paving the way to develop novel blockers, useful for the treatment of channelopathies such as neonatal diabetes (Remedi et al., 2017) or Cantú syndrome (Brownstein et al., 2013; Cooper et al., 2014). Further, our study provides insights into how RSG might exert its cardiovascular side effects, via interfering with its gating mechanism. We performed unbiased MD simulations on the microsecond time scale of the pore forming Kir6.1 model with RSG randomly placed in the solvent. After identification of a putative RSG binding site, we constructed dynamic pharmacophore models using the recently introduced common hits approach and screened for hits in *DrugBank*, which contains all drugs available on the market. Functional testing confirmed three new high affinity blockers, with different chemical scaffolds (see Figure 5). The identified compounds (see Table 2) provide an important first starting point for developing novel therapies for rare diseases such as CS. Taken together this study provides novel insights into the structural basis of Kir6.x channel block and may have broader implications for the molecular pharmacology of Kir6 channels in general.

## Methods

### Homology modelling

At the beginning of this study, no atomic resolution structure of a KATP channel was available. Thus, a Kir6.1 homology model in the closed state was built using the crystal structure of Kir3.2 (PDB code: 3SYA (Whorton and MacKinnon, 2011), 2.98 Å resolution) as template with the program Modeller9.11 (Martí-Renom et al., 2000). The sequence identity between Kir6.1 and Kir3.2 is 48.36%. The sequence alignments can be found in Supplementary Figure 1A. Comparison of the Kir3.2 template with recent available Kir6.2 structures reveal that the structures are highly similar with RMSD values below 1 Å (comparing the transmembrane domains). The structural alignments, generated with the Swiss-pdb-viewer (Guex and Peitsch, 1997) are shown in Supplementary Figure 1B.

### MD simulations

MD simulations were performed using Gromacs5.1 (Abraham et al., 2015) and the Amber99sb force field (Hornak et al., 2006). The Kir6.1 protein was embedded into the palmitoyloleoylphosphatidylcholine (POPC) lipid bilayer with four PIP_2_ molecules bound to the channel, as described previously (Lee et al., 2016). PIP_2_ was parameterized using the Hartree-Fock geometry optimization with the 6-31G* basis set (Frisch et al., 2013). POPC parameters were taken from Berger lipids parameters (Berger et al., 1997). The system was solvated using the SPCE water model (Berendsen et al., 1987; Kusalik and Svishchev, 1994) and 150 mM KCl were added to the solvent. To keep the selectivity filter stable, five K^+^ ions were placed at sites S0 to S4. The force field parameters of the ligand were generated and optimized with Gaussian09 (HF/6-31G* basis set) and antechamber (Wang et al., 2004, 2006). Ten R-form and ten S-form ligands were randomly placed in the solvent of the system. The algorithm to integrate Newton’s equation of motion was leap-frog, with a time step of 2 fs. The LINCS algorithm (Hess et al., 1997) was used to constrain all bonds. The cutoff-scheme for neighbor searching used Verlet (Verlet, 1967) within 1 nm and updated the list every 10 fs. The electrostatics and VdW interactions were measured with the particle-mesh Ewald (PME) method (Darden et al., 1993), using a cut-off of 1 nm and Fourier spacing of 0.16 nm. Temperature coupling used the V-rescale method (Bussi et al., 2007) at a reference temperature of 310 K and time constant 0.1 ps. The pressure was kept constant at 1 bar by using the Parrinello-Rahman barostat algorithm (Parrinello and Rahman, 1981) with a coupling constant of 2 ps. The system was minimized with the steepest descent algorithm, followed by a 6 ns equilibration simulation. 1.5 μs unbiased MD simulations were performed to detect the ligand binding sites. Additionally, 200 ns MD simulations were run from the best docking pose of the ligand.

### Docking

RSG was docked at the putative binding site identified from unbiased MD simulations, using the program Gold4.0.1 (Jones et al., 1997). The binding sites identified in the 1.5 μs free MD simulations were used as starting point and the radius was set to 20 Å. 100,000 operations of the GOLD genetic algorithm were used to dock the compounds with the ChemPLP scoring function.

### Umbrella sampling (US)

In order to estimate the ligand binding affinity, we performed US at each binding site. Ligands were firstly pulled into the solvent using the pull code in GROMACS by applying a harmonic biasing force between the center of mass (COM) of ligand and the COM of binding site (defined by residues within 5 Å of the ligand). The initial systems were taken from the last frames of the MD simulations for binding sites A_ref, B and C. To ensure that the ligands were pulled along the reaction coordinates fully into the solvent area, a harmonic force of 1,000 kJ/(mol·nm^2^) was applied for most of the pulling simulations. In cases, where the ligand displayed high mobility during the pulling trajectories, the harmonic force was increased to 2,000 kJ/(mol·nm^2^). Starting configurations for US were chosen from the pulling trajectories by taking steps every 0.1 nm along the reaction coordinates. Several intermediate windows were added if the adjacent US windows did not overlap sufficiently. Harmonic forces of 500, 1,000, 2,000 or 3,000 kJ/(mol·nm^2^) were applied to restrict the ligands during US sampling. For each window, a 10 ns simulation was performed, excluding the first 1 ns as equilibration. In total, 242 windows were simulated. Thus, in total, 2.42 μs simulations were performed to obtain good US window overlaps (Supplementary Figure 4). The potential of mean forces (PMF) were calculated by using weighted histogram analysis method (WHAM) (Hub et al., 2010) and the statistical errors were estimated by 100 times bootstrap analysis (Efron, 1979). A more detailed description about the US method can be found in the Supplementary Methods.

### Pharmacophore modelling

The recent published Common Hits Approach (CHA) (Wieder et al., 2017) was applied to construct dynamic pharmacophore models and to generate a hit-list by virtual screening in *DrugBank* (Law et al., 2014). The CHA is implemented by *LigandScout 4.10* (Wolber and Langer, 2005).

5,000 snapshots were extracted from the last 100 ns MD simulation of RSG at binding site A_ref and used as input for the CHA. For each snapshot, a pharmacophore model was built by considering the ligand interactions with protein and lipids. Water molecules were discarded during the pharmacophore generation. Pharmacophore features (mainly including hydrophobic interactions, hydrogen bond donor/acceptor, aromatic ring, ionizable area etc.) and constrains were defined as described in detail in the LigandScout user manual (LigandScout user manual, 2010).

Representative pharmacophore models were obtained by merging all identical features, extracted from the 5,000 frames. In the end, five representative pharmacophore models were used for virtual screening against *DrugBank4.0* (Law et al., 2014) (see Supplementary Figure 3). The molecules in *DrugBank* were prepared as libraries for virtual screening using the *LigandScout* command line tool *idbgen*. Conformers for each molecule in the database were generated using the icon best option in *idbgen*; this option produces a maximum number of 200 conformations for each molecule processed. The CHA produced a ranked hit-list for the binding site. The approved drugs that fits at least two of the five representative pharmacophore models were proposed in the final hit-list shown in Table 2.

### Interaction map

The interactions of RSG and N-RSG at site A_ref (only protein and PIP_2_ were considered) over the 200 ns MD simulations were analyzed and quantified by interaction maps, which were generated by the python package *matplotlib* (Hunter, 2007) and the chemoinformatic toolkit *CDPkit* (Seidel and Langer, 2017). Interactions were analyzed by generating a structure-based pharmacophore model at every saved frame of the MD trajectories and subsequently analyzing the frequency of the individual features. The interaction types were defined and described as pharmacophore features in the LigandScout user manual, including hydrophobic (H), hydrogen bond (HB) acceptor/donor, positive ionizable (PI) and aromatic (AR) features. The ligands were decomposed into five areas (Figure 4A and Supplementary Figure 2): the pyridine (Ring A), the benzene (Ring B), the thiazolidinedione (Ring C), the linker connecting pyridine and benzene (Linker D), and the linker connecting benzene and thiazolidinedione (Linker E). The frequencies of interactions observed were numbered and colored in the interaction map.

### Electrophysiology

Inside-out patch clamp electrophysiology was performed as described previously (Harakalova et al., 2012). In short, HEK293T cells were cultured on 10 mm glass coverslips and transfected with 0.16 μg of rat pCMV6-Kir6.2, 0.16 μg of rat pCMV6-SUR2A and 0.08 μg of pEGFP1 expression constructs. Measurements were performed using an AxoPatch 200B amplifier controlled by pClamp 9 software (Molecular Devices) at 22 °C using a ramp protocol ranging from −100 to +100 mV in 5 s from a holding potential of −40 mV. The sampling rate was 50 kHz, filter frequency was 2 kHz. Bath solution contained 131 mM KCl, 1 mM EGTA, 7.2 mM K_2_HPO_4_, 2.8 mM KH_2_PO_4_, 1 mM MgCl_2_, pH 7.20/KOH. The pipette solution contained 145 mM KCl, 1 mM CaCl_2_, 1 mM MgCl_2_, 5 mM HEPES, pH 7.40/KOH. Pipette resistance was 1.5-3 MΩ. Data were not corrected for rundown, which was less than 10% at 10 minutes. All measurements were performed within a timeframe of 8-10 minutes. Fractional block at −80 and +50 mV was determined by dividing current levels obtained with test compound containing solutions by current levels of control traces recorded in the absence of test compound.

Betaxolol (Sigma-Aldrich, St. Louis MO, USA) and Ritodrine (Sigma-Aldrich) were dissolved in H_2_O at 100 mM. Travoprost (MedChemExpress, Monmouth Junction, NJ, USA) was dissolved in DMSO at 10 mM. Test compounds were diluted in bath solution at the indicated concentrations before the start of measurements.

## Author contributions

XC, MW, AG and MJCH performed research, ASW and MvdH designed the study, XC, MW, EMZP, AG, MJCH, TL, MvdH and ASW analyzed data, ASW, MW, MvdH and XC wrote the paper. All authors reviewed the manuscript.

## Competing interests

The authors declare no competing interests.

## Funding

This work was supported by the Austrian Science Fund (FWF; http://www.fwf.ac.at). ASW and XC are supported by FWF grant I2101 (E-RARE 2). ASW, XC and EMZP are supported the doctoral program “Molecular drug targets” W1232 (FWF). MvdH and MJCH are supported by the E-Rare 2 Joint Transnational CantuTreat program.

## Acknowledgements

The computational results presented have been achieved in part using the Vienna Scientific Cluster (VSC).

