## supplementary material for "Computational identification of novel Kir6 channel inhibitors"

### Supplementary Methods

#### Potential of mean force (PMF) and umbrella sampling

PMF $\mathcal{W}(\xi)$ describes the free energy profiles of the system along reaction coordinates $\xi$, which can be defined as a distance, an angle, RMSD or more complicated functions. In our case, PMF describes the free energy changes of a ligand pathway, moving from the binding site to the solvent. The mathematical expression of PMF can be found in Roux, 1995 (Roux, 1995): The PMF $\mathcal{W}(\xi)$ can be defined from the average distribution function $\langle\rho(\xi)\rangle$ in equation (1):

$$\mathcal{W}(\xi) = \mathcal{W}(\xi^*) - k_B T ln\left[\frac{\langle\rho(\xi)\rangle}{\langle\rho(\xi^*)\rangle}\right] \quad (1)$$

where $\mathcal{W}(\xi^*)$ and $\langle\rho(\xi^*)\rangle$ are arbitrary reference values, $k_B$ is the Boltzmann constant and $T$ is the temperature. The average distribution function $\langle\rho(\xi)\rangle$ along the coordinate $\xi$ is obtained from a Boltzmann weighted average. In case an energy barrier present, the average distribution function cannot be computed due to the lack of sampling, non-Boltzmann sampling so called umbrella sampling (US) method can be applied to obtain PMF.

With US technique, the reaction coordinates space is divided into a series of windows. For each window, simulations are run with applying a harmonic biasing potential to the system, to ensure that high-energy regions are sufficiently sampled. The windows are well sampled when the distribution function at two adjective windows have good overlap (see histograms in Supplementary Figure S4, below). An unbiased PMF $\mathcal{W}_i(\xi)$ for each window $i$ is recovered from the biased simulations following equation (2):

$$\mathcal{W}_i(\xi) = \mathcal{W}(\xi^*) - k_B T ln\left[\frac{\langle\rho(\xi)\rangle_i^{biased}}{\langle\rho(\xi^*)\rangle}\right] - w_i(\xi) + F_i \quad (2)$$

where $\mathcal{W}(\xi^*)$ and $\langle\rho(\xi^*)\rangle$ are arbitrary reference values, $k_B$ is the Boltzmann constant and $T$ is the temperature, $\langle\rho(\xi)\rangle_i^{biased}$ is the average of biased distribution function of each window $i$ , $w_i(\xi)$ is the biasing harmonic potential applied to restrain the system, $F_i$ is undetermined constant that represents the free energy associated with $w_i(\xi)$.

The unbiased PMF $\mathcal{W}(\xi)$ of all windows and the constants $F_i$ can be computed by the weighted histogram analysis method (WHAM) (Kumar et al., 1992).

### Supplementary Figures

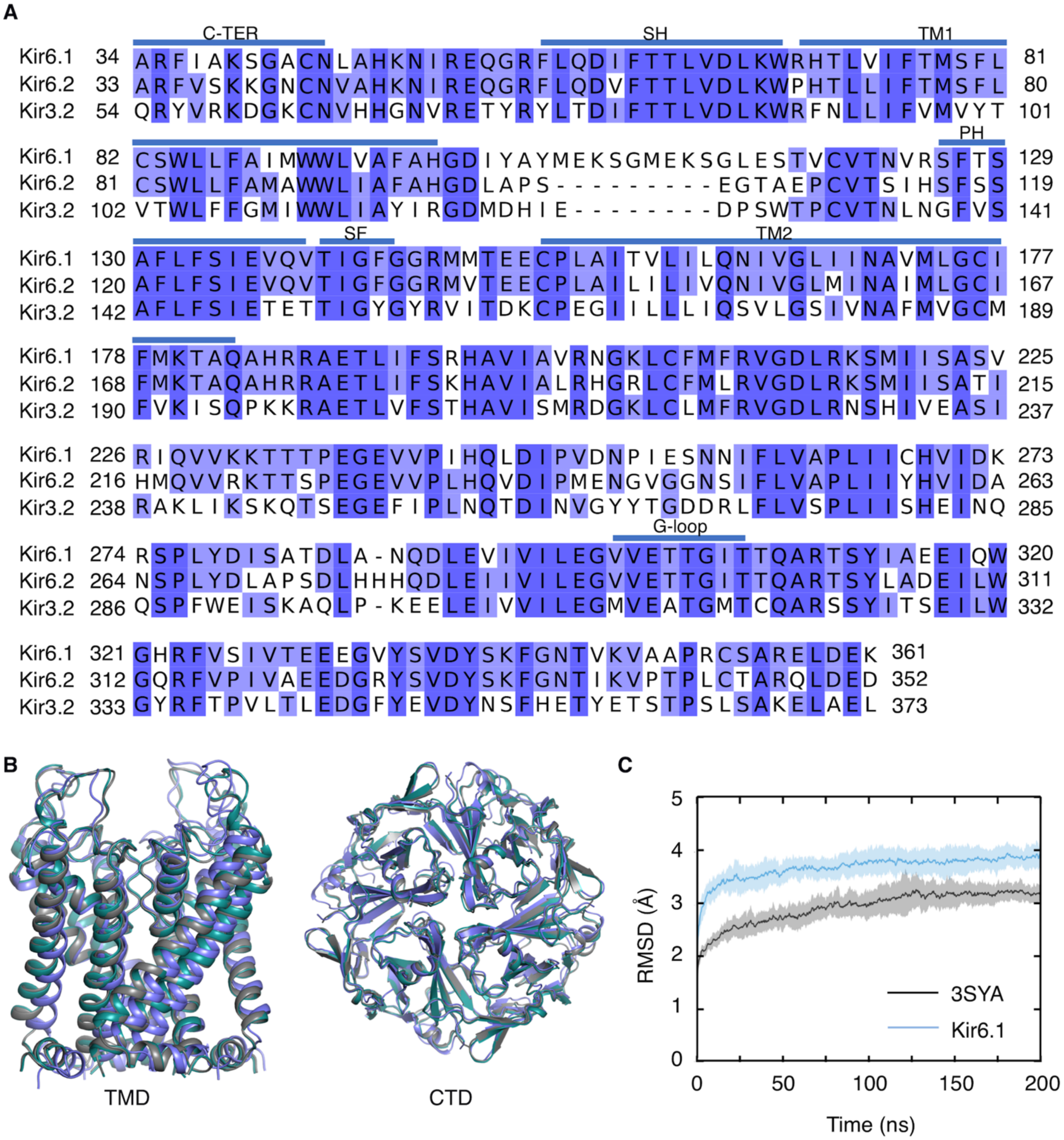


**Supplementary Figure 1. Kir6.1 modelling. (A)** Sequence alignment of human Kir6.1, Kir6.2 and Kir3.2 channels. **(B)** Cartoon representation of structural alignment about the TMD (side view) and the CTD (top view) of the Kir6.1 model in green, with the Kir3.2 template in grey (PDB code: 3SYA) and Kir6.2 in purple (PDB code: 6BAA). The RMSDs of structure alignments for the TMD and CTD part respectively are: 0.56 Å and 0.58 Å between Kir6.1 and Kir3.2, 0.86 Å and 1.34 Å between Kir6.1 and Kir6.2, and 0.92 Å and 1.02 Å between Kir6.2 and Kir3.2. **(C)** The average backbone RMSDs of the Kir6.1 model and the Kir3.2 template from three independent 200 ns MD simulations, show proteins are stable.

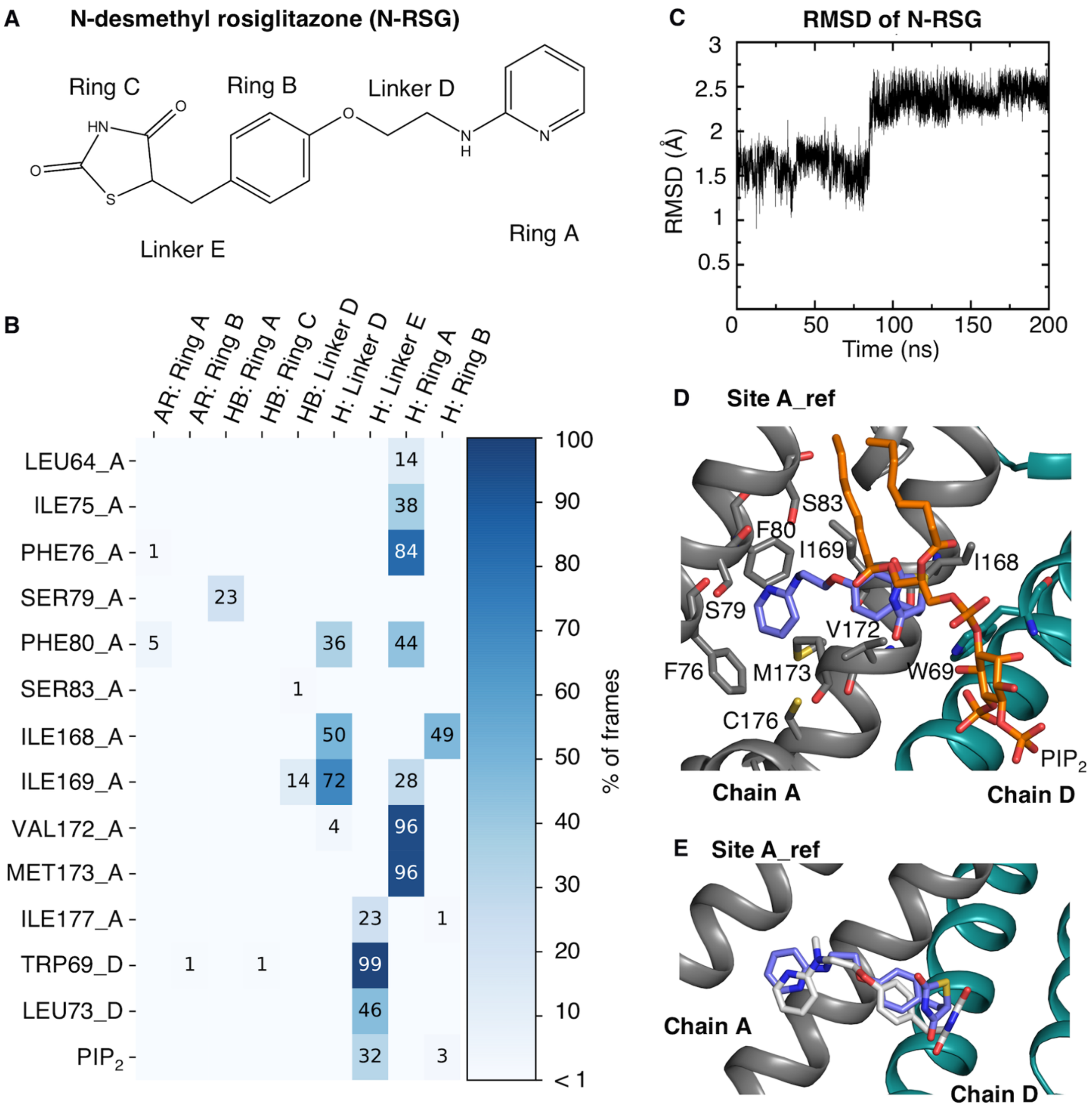


**Supplementary Figure 2. N-RSG interaction with the protein and $PIP_2$ at binding site A_ref.** **(A)** Molecular structure of N-RSG including the denotation corresponding to the interaction map. **(B)** Interaction map of N-RSG with protein and $PIP_2$ during 200 ns MD simulation. The matrix is coloured and numbered by the percentage of frames, in which the interactions were observed: aromatic (AR), hydrophobic (H), hydrogen bond (HB). The residues in the Kir6.1 are named by the corresponding amino acid with its residue number and chain ID (A or D). **(C)** RMSD plot of N-RSG at binding site A_ref during a 200 ns MD simulation. **(D)** Best PMF energy pose: Kir6.1 is represented as cartoon with the two neighbouring subunits coloured in grey and green, respectively. N-RSG (purple), the surrounding residues within 3.5 Å, and the $PIP_2$ (orange) are shown as stick. **(E)** Best PMF energy poses of RSG (white) and N-RSG (purple) superposed at binding site A_ref.

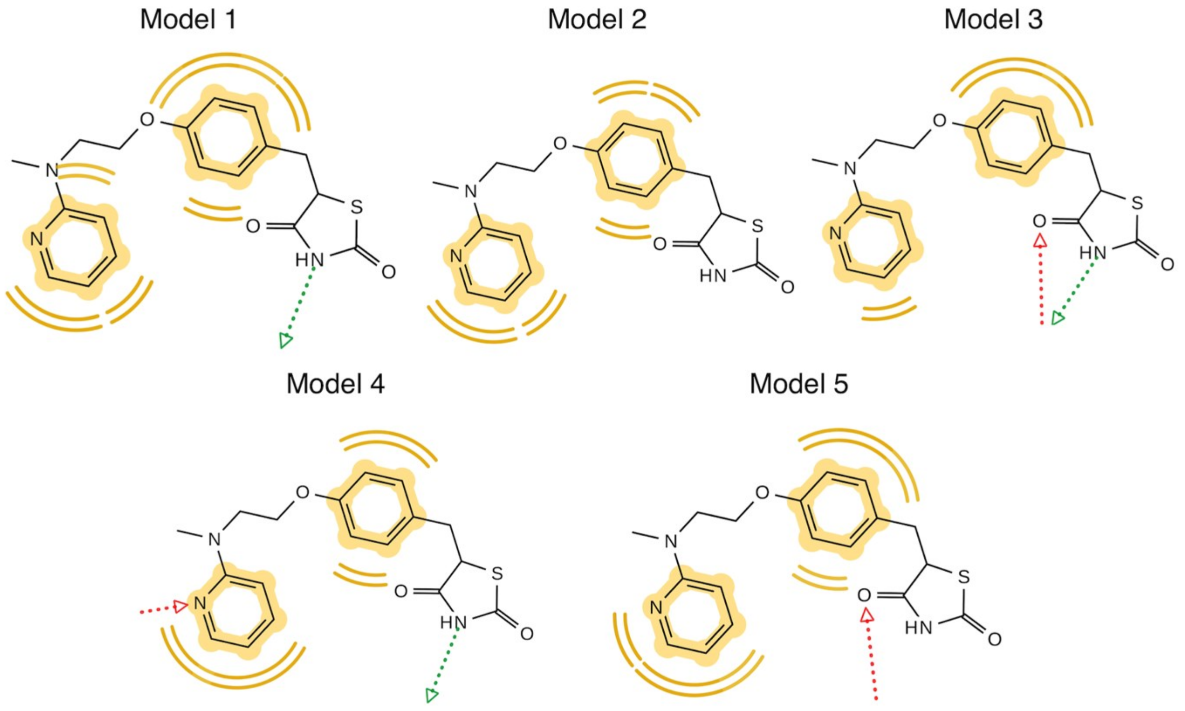


**Supplementary Figure 3. Dynamic pharmacophore models of RSG derived from MD simulation at binding site A_ref.** The yellow regions represent hydrophobic features. Red arrows represent hydrogen bond acceptors (HBA) and green arrows represent hydrogen bond donors (HBD).

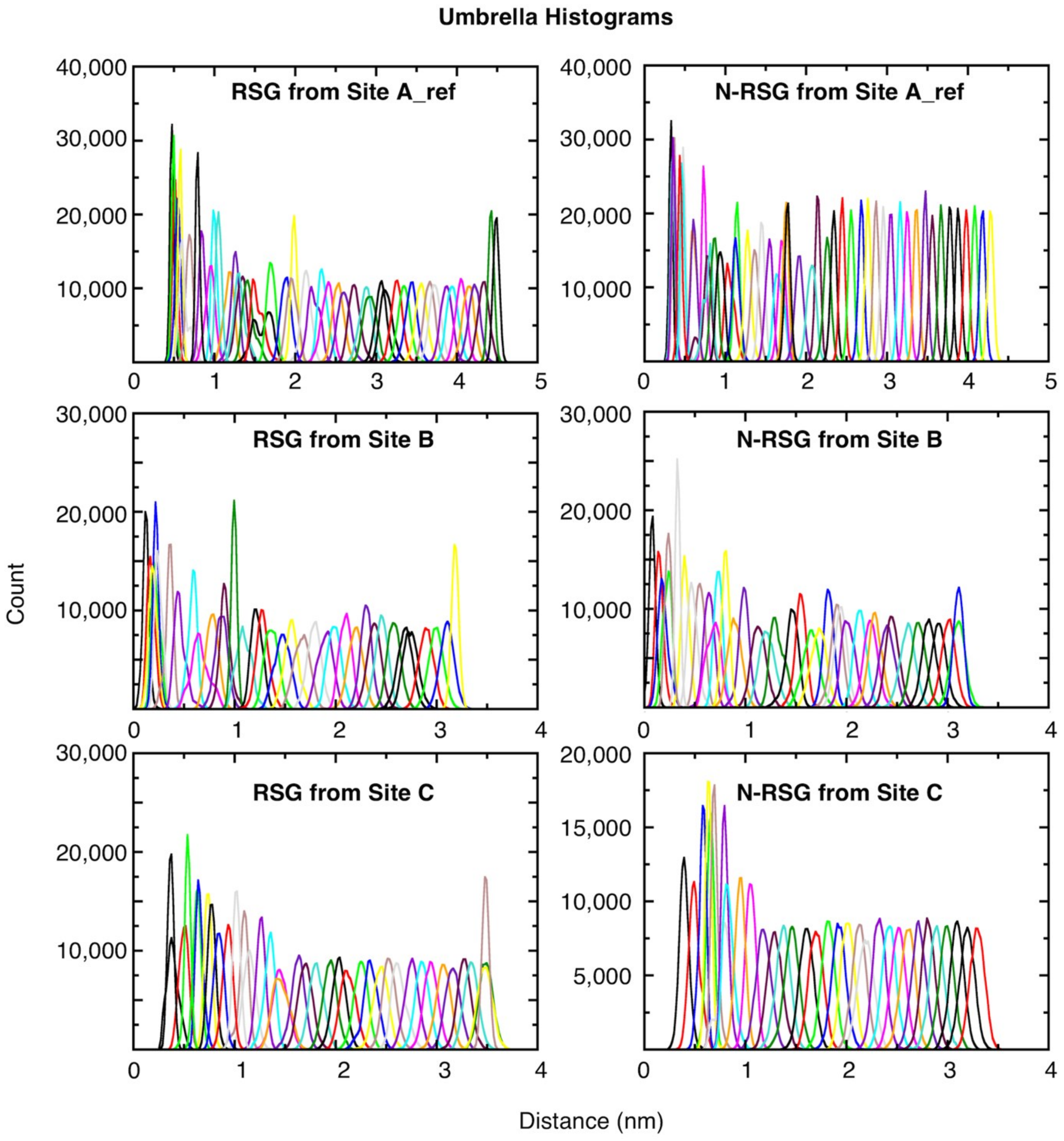


**Supplementary Figure 4. US histograms of RSG and N-RSG from the three distinctive binding sites.** Each plot represents the distribution function of a window, which shows good overlap. In total, 242 windows were generated with 10 ns simulation performed per window.

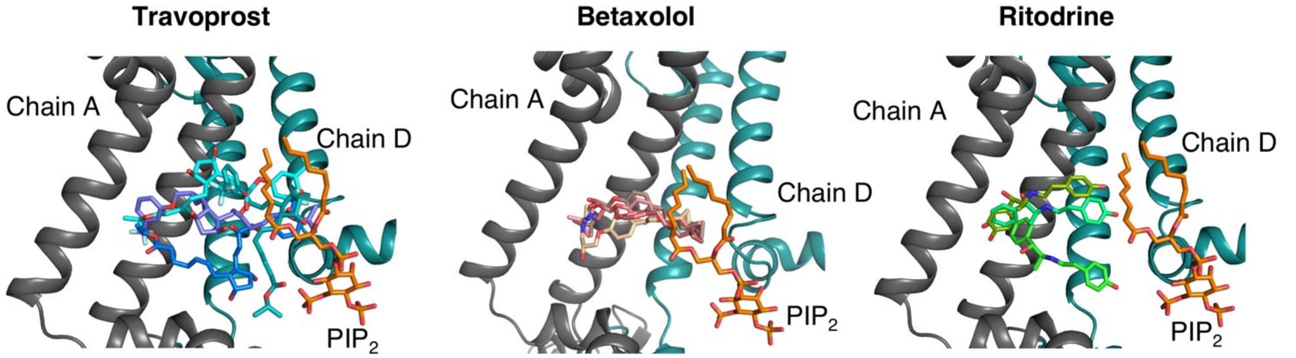


**Supplementary Figure S5. Docking poses of Travoprost, Betaxolol and Ritodrine at Site A_ref.** Kir6.1 is represented as cartoon with the two neighbouring subunits coloured in grey and green, respectively. The $PIP_2$ are shown as stick in orange. Docking poses of three ligands are represented as sticks at the binding site A_ref by different colours.

**Supplementary Tables**

**Supplementary Table S1. Indication, description and relevant side effect of the top ranked hit-list compounds at binding site A_ref.** All information is collected from *Sider* database (Kuhn et al., 2010, 2016) and *Drugbank* (Law et al., 2014).

| DrugBank ID | Generic name | Indication / Description | Relevant side effect |
|---|---|---|---|
| DB08907 | Canagliflozin | Anti-diabetic (sodium-glucose transport protein inhibitor) | Fungal genital infection funga, Urinary tract infection, Nausea |
| DB01095 | Fluvastatin | Antilipemic (inhibits HMG-CoA reductase) | Myotoxicity |
| DB09351 | Levobetaxolol | Glaucoma or ocular hypertension (selective beta-1-adrenergic receptor antagonist) | Burning or stinging in eye drainage from the eye redness; Swelling, and/or itching of eye and eyelid |
| DB00917 | Dinoprostone | Cervix preparation and induction for labour | Foetal heart rate; Foetal distress syndrome; Uterine |

| | | | |
|---|---|---|---|
| | | (naturally occurring prostaglandin E2, PGE2) | hypertonus; Failed induction of labour |
| DB00195 | Betaxolol | Glaucoma or ocular hypertension (cardioselective beta-1-adrenergic antagonist) | Fatigue; Dyspepsia; Headache; Dizziness; Arthralgia; Myalgia; Insomnia; Diarrhoea |
| DB00841 | Dobutamine | Cardiac stimulant after myocardial infarction or open heart surgery (beta-1 agonist) | Angina pectoris; Chest pain; Headache |
| DB00287 | Travoprost | Glaucoma or ocular hypertension (ophthalmic solution, synthetic prostaglandin F2alpha analog) | Hyperaemia; Iris hyperpigmentation; Eye disorder; Ocular hyperaemia; Eye pruritus; Eye pain and irritation; Ocular discomfort |
| DB00938 | Salmeterol | Asthma and chronic obstructive pulmonary disease (long-acting beta2-adrenergic receptor agonist) | Headache; Musculoskeletal pain; Pallor; Sinus congestion; Hypersensitivity Tremor; Palpitations |
| DB00179 | Masoprocol | Actinic keratoses (precancerous skin growths that can become malignant if left untreated; lipoxygenase inhibitor) | Antineoplastic |
| DB00867 | Ritodrine | Treatment and prophylaxis of premature labour (beta-2 adrenergic agonist) | Increase in heart rate; Arrhythmia |
| DB09198 | Lobeglitazone | Antidiabetic drug (peroxisome proliferator-activated receptor (PPAR) alpha and gamma) | Not approved to use |

| | | | |
|---|---|---|---|
| DB04855 | Dronedarone | Class III antiarrhythmic | Pulmonary fibrosis; Interstitial lung disease; Leukocytoclastic vasculitis |
| DB06817 | Raltegravir | Antiretroviral treatment of HIV infection (integrase inhibitor) | Malnutrition; Mental disorder |
| DB09570 | Ixazomib | Multiple myeloma (proteasome inhibitor) | Antineoplastic; Hepatotoxic |
| DB01346 | Quinidine barbiturate | - | Obsolete |
| DB00204 | Dofetilide | Class III antiarrhythmic (IKr inhibition) | Torsades de pointes |
| DB01240 | Epoprostenol | Primary pulmonary hypertension (prostaglandin I2, vasodilator, inhibition of platelet aggregation) | Headache; Hypotension; Nausea |
| DB00662 | Trimethobenzamide | Antiemetic (D2 receptor antagonist) | Parkinsonism; Tremors |
| DB08875 | Cabozantinib | Metastatic medullary thyroid cancer, renal cell carcinoma (non-specific tyrosine kinase inhibitor) | Gastrointestinal fistulas and perforations; Potentially fatal hemoptysis and gastrointestinal hemorrhage |
| DB09330 | Osimertinib | Non-small cell lung cancer (epidermal growth factor receptor tyrosine kinase inhibitor) | Interstitial lung disease |
